# Sirtuin 3 with rs11607019 regulates gene expression in 293 cells

**DOI:** 10.64898/2025.12.03.692027

**Authors:** Yue Chen, Chengyu Liu, Wei Ju, Shiqi Wang, Ying Hu, Yadong Wang

## Abstract

**Background:** SIRT3 is a NAD+ dependent deacetylase located in mitochondria and plays a critical role in various biological processes such as aging and cancer. Recent GWAS analyses have identified an intronic SNP, rs11607019, that significantly influences platelet function.

**Methods:** To investigate the genome-wide transcriptional effects of this variant, we performed site-directed mutagenesis experiments.

**Results:** Our results demonstrate that rs11607019 upregulates GABRE, COL12A1, PLEKHG4B, and SERF1B, while downregulating APOBEC3G in 293 cells, without cell tumor mutational burden karyotype phenotype.

**Conclusion:** These findings suggest that rs11607019 is a strong candidate locus for blood cell phenotyping and may warrant further investigation in relation to kidney-related diseases in latent populations.

## Background

Sirtuin 3 (SIRT3) is an NAD+-dependent deacetylase [1] predominantly localized in mitochondria. Recent studies by Diao et al. demonstrated that SIRT3 mitigates cellular senescence, in part through heterochromatin stabilization [2]. Besides its role in maintaining mitochondrial function and metabolic homeostasis [3], SIRT3 is involved in stress response and transcriptional regulation [4]. Additionally, SIRT3 has been involved in various pathologies, including acute kidney injury [5, 6] and cancer [7].

Recently, Chen et al. [8] conducted genome-wide association study (GWAS) of 15 hematological traits in 746,667 Individuals from 5 Global Populations. In the study, they meta-analyses the Trans-ethnic and Ancestry-Specific Blood-Cell clinically relevant complex human traits across diverse ancestral groups and identified rs11607019 to be associated with platelet phenotype in Trans-ethnic and Ancestry-Specific Blood Cell Trait. Another study genotyped 435 individuals with type 1 diabetes (T1D) using the Illumina Infinium Omni Express Exome-8 v1.4 array and conducted a mitochondrial GWAS (mitoGWAS) for BMI [9] . The analysis revealed an interaction between SIRT3 variant rs11607019 and MT-ND2 variant rs28357980, both of which are implicated in mitochondrial function and energy metabolism. These variants were associated with BMI in young women with T1D undergoing insulin treatment (MAF = 10.3%, p-value = 0.056). To date, no pathological phenotypes have been linked to rs11607019 (Table S1). Collectively, these findings indicate that rs11607019 is a functionally relevant variant warranting additional research.

To further investigate the functional role of rs11607019, we induced mutation and analyzed its mediated gene expression in conventional human cell models. Our results demonstrated that rs11607019 regulates the expression of GABRE, COL12A1, PLEKHG4B, SERF1B, and APOBEC3G. A literature review revealed a strong association between these genes and platelet volume. However, the precise relationship between this locus and blood cell phenotypes remains inconclusive, warranting further exploration across diverse tissues and populations to elucidate potential health disparities.

## Methods

### Plasmid, cell and genotype

The HEK293 cell line was obtained from the ATCC and maintained at UBIGENE (Guangzhou, China). The ABEmax plasmid (Addgene, Catalog #112095) was used for base editing. The single-guide RNA (sgRNA) targeting the desired genomic locus (sequence: 5′-TGAGGAGTAGTCCATTGCATGGG-3′) and PCR genotyping primers (Forward: *5′-CGTGGAGCGCATTCTAATATCTGTG-3′*, Reverse: *5′-CATCTATTCACTTAAGGTCGTTCC-3′*) were synthesized by Sangon Biotech (Shanghai, China).

### RNA-seq

RNA-seq was performed by BGI (Shenzhen, China) using the DNBseq platform with read length of SE50 and DNBSEQ Eukaryotic Strand-specific mRNA library.

### Sequencing data process

Raw sequencing reads were aligned to the hg38 reference genome using HISAT2. The aligned reads were then sorted and indexed using SAMtools, and read counts were generated with HTSeq-count.

### Differential expression analysis

Differential gene expression analysis was conducted using the DESeq2 package in R with default parameters. Genes with significant expression changes were identified based on an adjusted p-value (padj, Benjamini-Hochberg FDR correction) and log2 fold change (log2FC) thresholds.

### G-banding karyotype

293 cells were collected and ongoing trypsin Giemsa staining [10] . Cell karyotypes were visualized with Zeiss Imager-Z2 microscope, randomly acquired at least 60 cell images, analysis with Ikaros karyotyping system.

## Results

### Selection of the rs11607019 Locus

The rs11607019 variant is located within the intronic region of SIRT3 on chromosome 11 (GRCh38 position: 11:234,349), with T/C alleles (forward strand) and a minor allele frequency (MAF) of 0.09 (Figure 1A, B). This locus exhibits strong statistical significance (*P*-value from GWAS Catalog) in association with mean platelet volume (Figure 1C). Furthermore, analysis of NCBI expression data revealed that SIRT3 is highly expressed in human kidney tissue compared to other tissues (Figure 1D).

**Figure 1.**
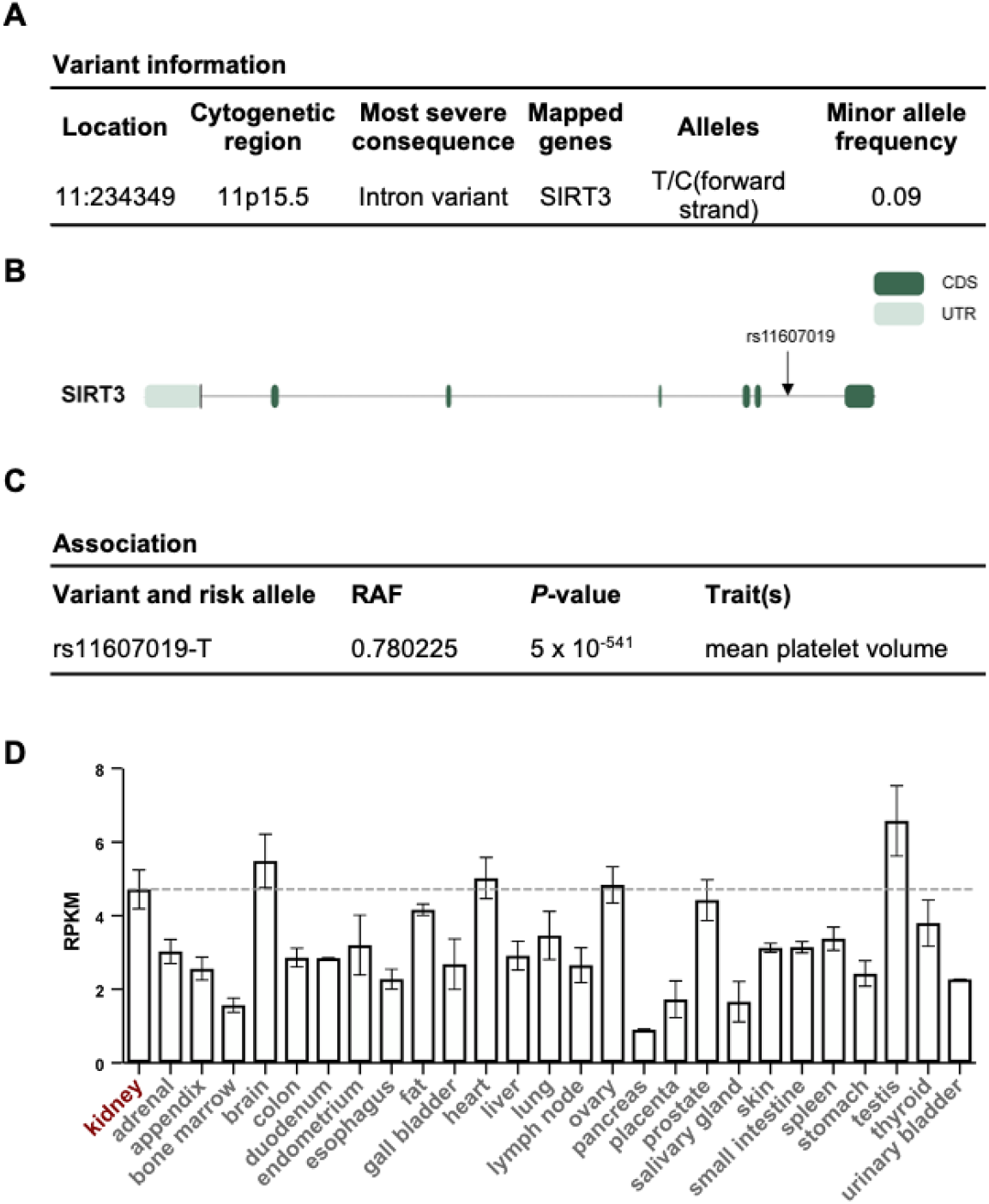
The overall information of rs11607019.

### Recombinant Vector Construction

The gRNA was designed using BE-Designer and synthesized as double-stranded DNA fragments. These fragments were ligated into BbsI-digested pUC57 expression vectors through overnight incubation at 16°C. The ligation products were then transformed into competent E. coli (30-50 μL) by heat shock (42°C for 90 seconds) following a 30-minute ice incubation. Transformed bacteria were recovered in antibiotic-free LB medium at 37°C for 30 minutes with shaking, then plated on LB agar containing 0.1% ampicillin. After 12-hour incubation at 37°C, single colonies were selected for overnight culture in liquid LB with 0.1% ampicillin (37°C, 12 hours). Plasmids were extracted using a commercial mini-prep kit and verified by sequencing (Figure 2).

**Figure 2.**
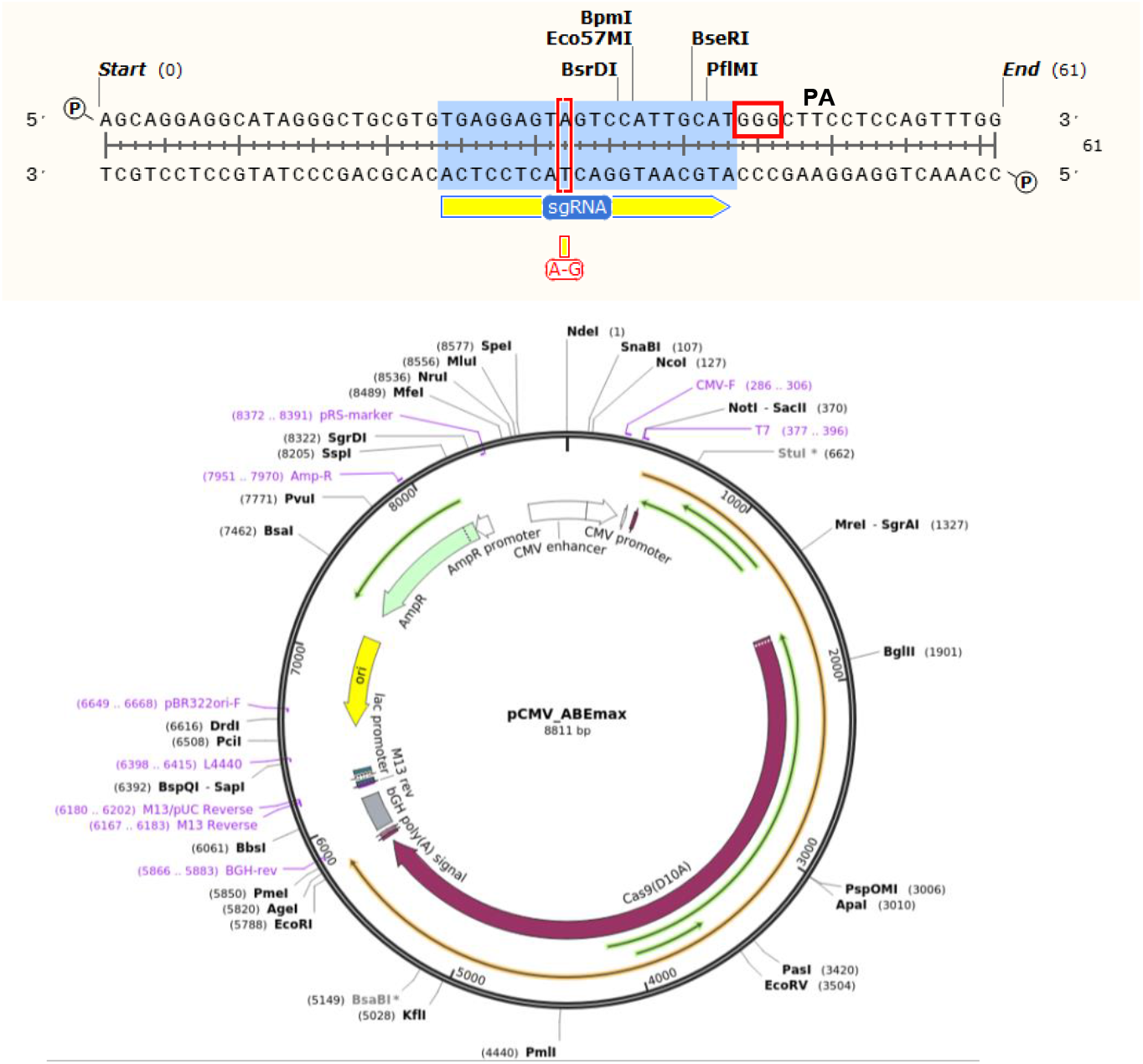
Piont mutate strategy.

### Generation and Characterization of Point-Mutated Cells

Using base editor technology [11], we successfully introduced the 11:234349T>C mutation (rs11607019) in 293 cells (Figure 3). Quantitative analysis revealed no significant alteration in SIRT3 expression in mutant cells compared to controls. However, differential expression analysis identified four significantly upregulated genes (GABRE, COL12A1, PLEKHG4B, and SERF1B) and one upregulated gene (APOBEC3G) (Table 1).

**Table 1.** Differential gene expression.

| Ensembl ID | Official Symbol | log2FoldChange | padj |
| --- | --- | --- | --- |
| ENSG00000142082 | SIRT3 | -0.266652293063418 | 0.901111045555107 |
| ENSG00000239713 | APOBEC3G | -2.30087844610027 | 1.26538572952605E-05 |
| ENSG00000102287 | GABRE | 4.54301309696295 | 1.30394737793688E-12 |
| ENSG00000111799 | COL12A1 | 2.2256409109775 | 2.96729982215507E-12 |
| ENSG00000153404 | PLEKHG4B | 2.02944179922927 | 0.0176947151716473 |
| ENSG00000205572 | SERF1B | 3.10854473521247 | 0.0376558061252757 |

**Figure 3.**
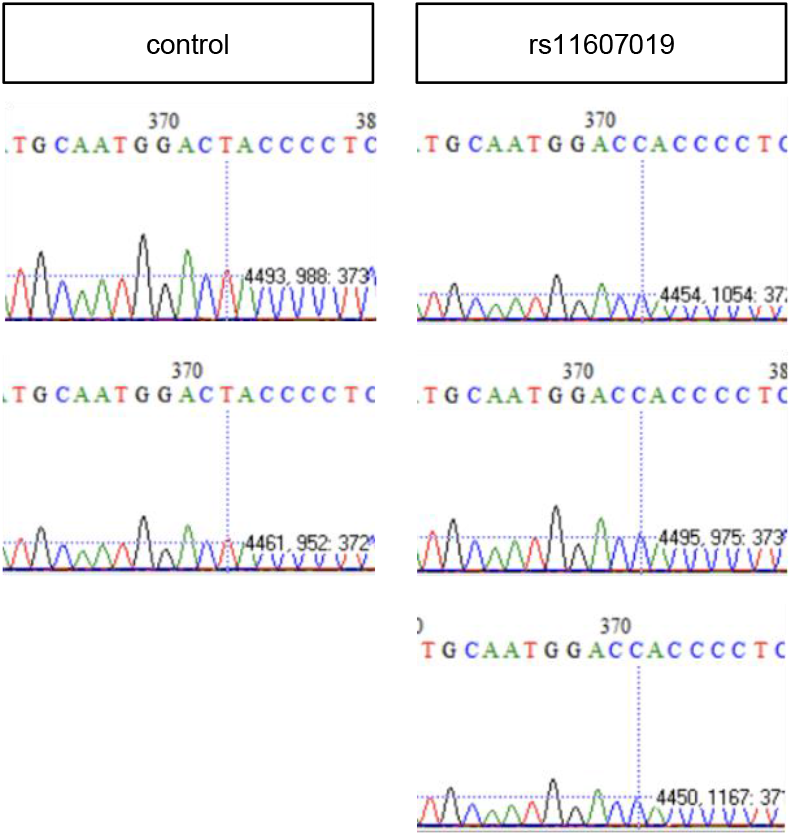
Genotyphing of 293 cells with/without rs11607019.

### The G-band pattern of rs11607019 293 cells

In our study, G-band staining was used for HEK293 cells chromosome analysis. As shown in Figure 4, after cell chromosome number rearranged to 46, chromosomes 1, 2, 5, 7, 10, 15, 18 were consistently diploid in all.

**Figure 4.**
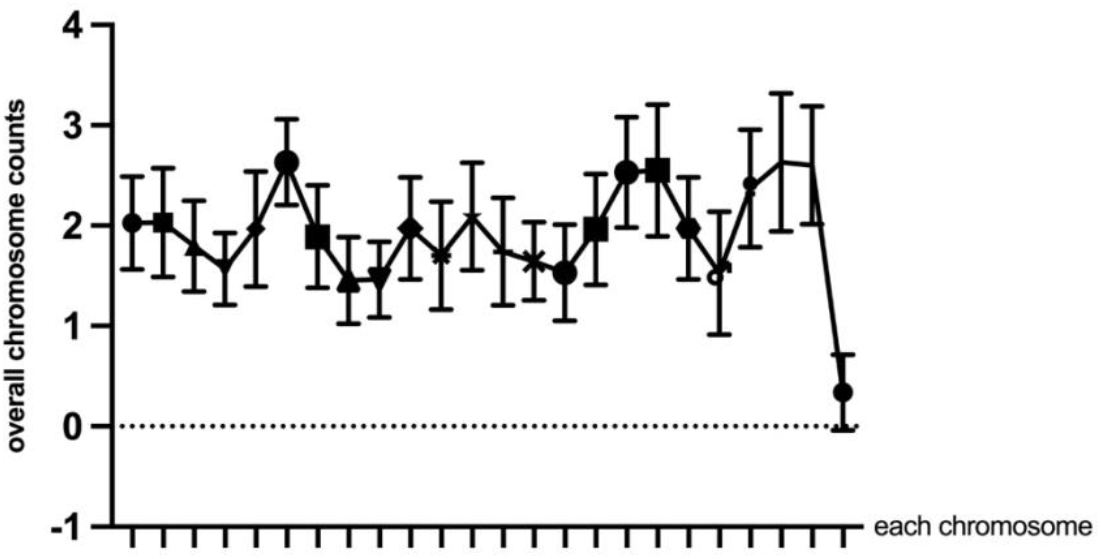
Karyotype of 293 cells with Sirt3 point mutation

## Discussion

Our study reveals that the rs11607019 variant modulates the expression of several functionally important genes. Notably, we identified four significantly upregulated genes (SERF1B, GABRE, PLEKHG4B, and COL12A1) and one moderately upregulated gene (APOBEC3G) in cells carrying this variant.

SERF1B, located within a 500kb inverted duplicated region containing repetitive elements, has been implicated in age-related proteotoxicity regulation [12]. GABRE encodes the γ-aminobutyric acid type A receptor ε subunit [13], which forms chloride channels critical for fast inhibitory synaptic transmission in the central nervous system. PLEKHG4B, a guanine nucleotide exchange factor with preferential expression in kidney, thyroid and testis (NCBI), may contribute to renal cellular signaling [14].

Of particular interest is APOBEC3G, which not only inhibits HIV-1 infectivity [15] but also promotes cancer mutagenesis in bladder cancer [16]. COL12A1, encoding a collagen XII α chain [17], mediates collagen I fibril-matrix interactions and facilitates breast cancer metastasis [18]. The collagen gene (COL) family has established roles in platelet function; for instance, COL8A1 modulates vascular stiffness through platelet-derived extracellular vesicles in arterial injury models [19]. These findings align with previous reports that rs11607019 influences mean platelet volume [8], suggesting potential mechanisms through which this variant may affect hematological traits.

## Conclusions

While mean platelet volume (MPV) is influenced by multiple factors including platelet count, demographic characteristics, and genetic background, our study provides the first experimental validation of rs11607019’s functional impact in human cells. These findings expand our understanding of genetic loci contributing to interindividual variation in blood cell traits, with potential implications for hematological disorders and aging-related processes [20]. Future studies should explore the precise mechanisms linking these transcriptional changes to platelet physiology and disease susceptibility.

## Declarations

### Ethics approval and consent to participate

Not applicable.

### Consent for publication

All authors read and approved the final manuscript.

### Availability of data and material

The datasets generated and analysed during the current study are available in the NCBI Gene Expression Omnibus (GEO) under accession number GSE299443.

### Authors’ contributions

YC, CL, WJ and SW conceived the idea, designed the research, acquired the funding, and contributed reagents. WJ and SW performed the experiments. All authors participated in data analysis and initial draft preparation. YH and YW finalized the manuscript.

## Acknowledgements

ABEmax plasmids is gifted from Prof. Zhanjun Li (Jilin University).

## Authors’ information

Not applicable.

